# Norepinephrine activates β1-adrenergic receptors localized to the inner nuclear membrane in cortical astrocytes

**DOI:** 10.1101/2021.06.24.446972

**Authors:** Kelsey C. Benton, Daniel S. Wheeler, Beliz Kurtoglu, Mahshid Bagher Zadeh Ansari, Daniel P. Cibich, Dante A. Gonzalez, Matthew R. Herbst, Saema Khursheed, Rachel C. Knorr, Doug Lobner, Jenree G. Maglasang, Kayla E. Rohr, Analisa Taylor, Paul J. Witt, Paul J. Gasser

## Abstract

Studies in cardiomyocytes have established that adrenergic receptors, conventionally thought to initiate signaling events exclusively from the plasma membrane, can also localize to and signal from the nuclear membrane. Activation of these receptors by their endogenous cationic ligands requires transmembrane uptake mediated by organic cation transporter 3 (OCT3). We have demonstrated that OCT3 is densely localized to outer nuclear membranes in neurons and astrocytes, suggesting that nuclear adrenergic signaling is also present in the central nervous system. In this study, we examined the subcellular localization of β1-adrenergic receptors, their G-protein signaling partners, and catecholamine transporters in mouse astrocytes. We identified a population of β1-adrenergic receptors localized to astrocyte inner nuclear membranes. We demonstrated that key components of G_s_-mediated signaling are localized to the nuclear compartment and identified OCT3 and other catecholamine transporters localized to plasma and nuclear membranes. Treatment of astrocytes with norepinephrine induced rapid increases in nuclear PKA activity which were blocked by pretreatment with inhibitors of catecholamine transport. These data indicate that nuclear β1-adrenergic receptors are functionally coupled to G_s_-coupled signaling mediators and that their activation by norepinephrine requires transporter-mediated uptake. These receptors represent a powerful mechanism by which norepinephrine may alter astrocyte gene expression and brain function.

## INTRODUCTION

G-protein-coupled receptors mediate powerful influences of diverse ligands on cellular function through signal transduction pathways that alter the phosphorylation of proteins throughout the cell. The specific effect of an extracellular ligand on a given target cell depends not only on the number and class of receptors expressed by the cell, but also on the localization of receptors in relation to intracellular effectors. Recent studies have demonstrated that, in addition to the plasma membrane, G-protein-coupled adrenergic receptors may also be localized to, and activated at, intracellular membranes including Golgi apparatus and inner nuclear membranes, and that receptors in distinct cellular locations mediate distinct cellular responses to catecholamines (Boivin et al., 2006; Dahl et al., 2018; Irannejad et al., 2017; Vaniotis et al., 2011; Wang et al., 2020; Wu et al., 2014). These studies have also shown that access of norepinephrine, a cationic and therefore membrane-impermeable ligand, to adrenergic receptors in the inner nuclear and Golgi membranes is mediated in part by organic cation transporter 3 (OCT3), a high-capacity transporter for norepinephrine and other monoamines (Irannejad et al., 2017; Wang et al., 2020; Wright et al., 2008). In addition to its actions on peripheral targets, norepinephrine is a potent regulator of integrated central nervous system function through its actions on neuronal and glial targets. Adrenergic receptors localized to the nuclear membrane represent powerful mechanisms by which norepinephrine may regulate diverse aspects of cellular function. However, no studies have examined the localization of adrenergic receptors to endomembranes in neuronal or glial cells.

We recently demonstrated using immunoelectron microscopy that, in addition to plasma membrane sites, OCT3 is densely expressed in the outer nuclear membranes of astrocytes in the rat and mouse brain (Gasser et al., 2017). The presence of this catecholamine transporter in the outer nuclear membrane suggested that adrenergic receptors may be localized to inner nuclear membranes in astrocytes, and that these receptors may mediate previously described effects of norepinephrine on astrocyte gene expression and physiology. Here, we examined the subcellular localization of β1-adrenergic receptors (β1-AR), their signaling partners, and catecholamine transporters in mouse astrocytes. We confirmed the nuclear localization of OCT3 in primary astrocytes and identified a population of β1-ARs localized to astrocyte inner nuclear membranes. We demonstrated that key components of G_s_-mediated signaling are localized to the nuclear compartment and identified multiple catecholamine transporters in addition to OCT3 localized to plasma and nuclear membranes in astrocytes. Finally, we tested the hypothesis that activation of nuclear β1-ARs by norepinephrine requires catecholamine transporter-mediated uptake. Treatment of astrocytes with norepinephrine induced rapid increases in nuclear PKA activity which were blocked by pretreatment with inhibitors of catecholamine transport. These findings reveal a previously undescribed mechanism by which norepinephrine regulates nuclear PKA activity in astrocytes, and through which norepinephrine may exert powerful influences on astrocyte gene expression, contributing to important metabolic, neuroprotective, and immunomodulatory actions of norepinephrine in astrocytes.

## RESULTS

### OCT3 is localized to astrocyte nuclei

To confirm that OCT3 is localized to the nuclear envelope in cultured astrocytes as it is *in situ*, we used an antibody directed against a peptide in the large intracellular loop of mouse and rat OCT3 (Alpha Diagnostic International, cat# OCT31-A, RRID:AB_1622571; (Gorboulev et al., 2005)) in immunofluorescence and immunoblotting studies. Punctate OCT3-like immunoreactivity was observed over cell bodies and nuclei of nearly all cultured astrocytes (**Fig. 1A-C**), with higher density over nuclei than over cell bodies.

**Fig. 1.**
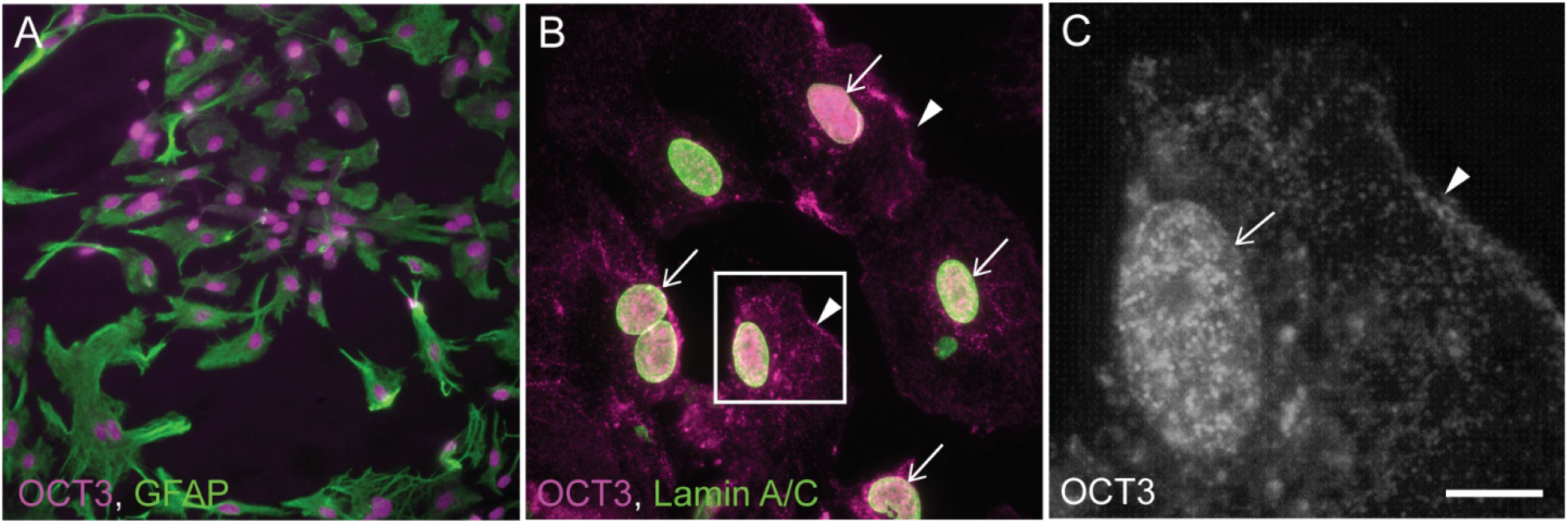
OCT3 is expressed in astrocyte nuclei. Fluorescence photomicrographs depicting OCT3 immunoreactivity (magenta in **A**, **B**; white in **C**); GFAP immunoreactivity (green in **A**only); and lamin A/C immunoreactivity (green in **B**only) in primary cultured mouse astrocytes. Box in (**B**) indicates area shown at higher magnification in (**C**). OCT3-like immunoreactivity is localized over apparent nuclei in all GFAP+ astrocytes (**A**). At higher magnification (**B, C**), OCT3-like immunoreactivity is observed over cell bodies (arrowheads in **B, C**) and nuclei (arrows in **B**, **C**). Images are representative of n = 5 independent experiments. Scale bar, 80 μm (**A**), 25 μm (**B**), 6.25 μm (**C**).

### β1-adrenergic receptor is localized to both plasma and inner nuclear membranes of astrocytes

To visualize the subcellular localization of β1-AR in astrocytes, we used two different commercially available antisera, each directed against a distinct epitope of the receptor, in immunofluorescence studies in cultured mouse astrocytes and frozen brain sections. One antibody (Thermo Fisher, cat# PA1-049, RRID:AB_2289444) is directed against an amino acid sequence in the C-terminal tail of mouse β1-AR which is an intracellular epitope of the plasma membrane-localized receptor. The second antibody (Alomone Labs, cat# AAR-023, RRID:AB_2340886) is directed against an amino acid sequence in the second extracellular loop of the plasma membrane-localized receptor. Nuclei were identified either by counterstaining with the DNA stain DAPI, or by immunolabeling with an antibody against the nuclear proteins lamins A and C.

The two β1-AR antibodies produced similar patterns of immunofluorescence in triton-permeabilized GFAP+ astrocytes (**Fig. 2A, B**). Punctate β1-AR-like immunoreactivity (β1-like ir) was diffusely distributed over astrocyte somata and densely localized over nuclei. Putative nuclear β1-like ir was reliably observed in nearly all cultured astrocytes. This pattern of staining was observed in 11 repeated assays with the C-terminal antibody and 7 repeated assays with the ECL2 antibody (each replicate used astrocyte cultures from distinct mouse litters). β1-AR-like-ir was not observed over nuclei or somata when immunofluorescence was conducted using primary antibodies which had been pre-adsorbed with an excess of the respective antigenic peptide (**Fig. 2C, D, G, H**).

**Fig. 2.**
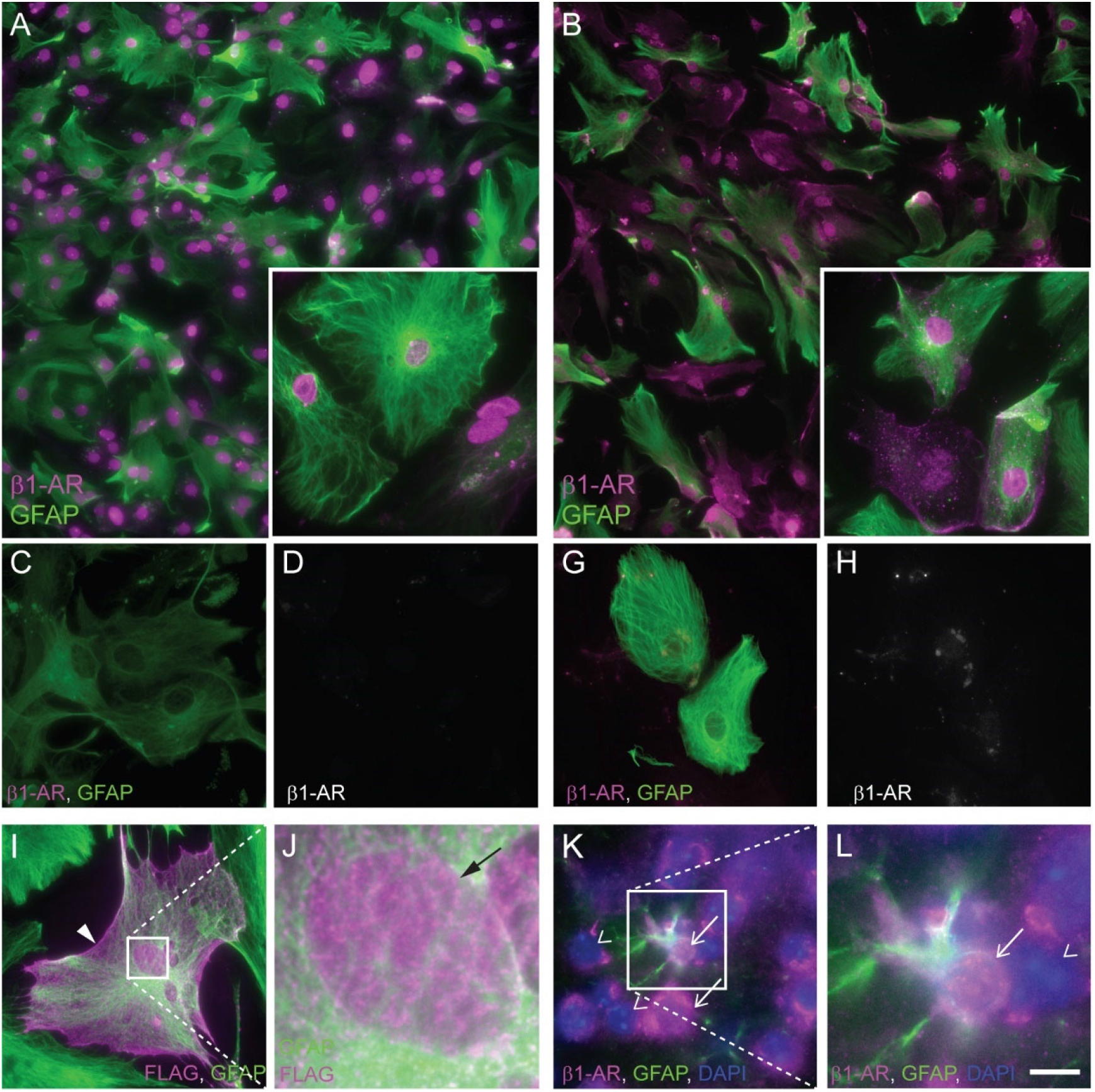
β1-adrenergic receptor localization in astrocytes. Fluorescence photomicrographs depicting β1-AR immunoreactivity in mouse primary astrocytes. β1-AR (magenta) was detected with an antibody against extracellular loop 2 (**A, C, D**) or the C-terminal tail (**B, G, H**) of the receptor. GFAP immunoreactivity is shown in green. Antibodies used in **C, D, G**, and **H**were pre-adsorbed with the immunizing peptide prior to application to cells. (**I**, **J**) Immunofluorescence of primary mouse astrocytes transfected with a FLAG-tagged human β1-AR and incubated with antibodies against FLAG peptide (magenta) and GFAP (green). Box in **I**indicates area shown at higher magnification in **J**. Arrowhead in **I**indicates cell body-localized FLAG (putative β1-AR) immunoreactivity. Arrow in **J**indicates nuclear FLAG (putative β1-AR) immunoreactivity. (**K, L**) Fluorescence photomicrographs depicting β1-AR (magenta) and GFAP (green) immunoreactivity in mouse prefrontal cortical tissue. DAPI labeling of DNA is depicted in blue. Arrows indicate β1-AR-immunoreactive nuclei. Arrowheads indicate β1-AR-immunonegative nuclei. Insets in A, B are higher magnification images from the same experiment. Images are representative of n ≥ 3 independent experiments (except for **C, D**, for which n=2 experiments). Scale bar, 62.5 μm (**A**, **B**), 25 μm (**C, D, G, H, I,**insets in **A, B**); 3.7 μm (**K**); 9.25 μm (**L**).

To confirm that the nuclear immunofluorescent signal represented β1-adrenergic receptor and not a cross-reactive protein, we transfected cultured mouse astrocytes with plasmids directing the expression of an N-terminal FLAG peptide-tagged human β1-AR (Tang et al., 1999), and conducted immunofluorescence using an anti-FLAG peptide antibody ~2 days after transfection. Anti-FLAG immunoreactivity was observed over nuclei and diffusely over cell bodies of transfected astrocytes (**Fig. 2I, J**). To determine whether nuclear β1-like ir is present in mouse cortical astrocytes *in situ*, we incubated fixed, frozen brain sections of mouse forebrain with the C-terminal-directed β1-AR antibody. β1-AR-like immunoreactivity was observed over both nuclei and somata in mouse prefrontal cortex (**Fig. 2K, L**). Identification of astrocyte nuclei is difficult in sectioned brain tissue using GFAP-immunoreactivity, as cortical astrocytes *in situ* express low levels of GFAP (Verkhratsky and Nedergaard, 2018) and because astrocyte processes extend three-dimensionally throughout the tissue, making it difficult to determine definitively whether a given nucleus belonged to a GFAP+ cell. However, nuclear β1-AR-like immunostaining was observed in a small number of nuclei that were obviously surrounded by GFAP-immunofluorescence.

The immunofluorescent signal observed over astrocyte nuclei could represent receptor localized to either inner or outer nuclear membrane, to plasma membrane overlying the nucleus, or to perinuclear endoplasmic reticulum. To determine whether the nuclear immunofluorescence originates from receptors localized to the inner or outer nuclear membranes, we conducted immunostaining using each of the two β1-AR antibodies on astrocytes permeabilized with either Triton X-100 or digitonin. Triton X-100, which solubilizes all cellular membranes, allows free access of antibodies to all cellular compartments. Digitonin at the concentration used here preferentially permeabilizes cholesterol-rich membranes like the plasma membrane, leaving inner and outer nuclear membranes mainly intact (Adam et al., 1990). Thus, in digitonin-permeabilized cells, antibodies would not be able to bind antigens in the nuclear lumen or the perinuclear space (between outer and inner nuclear membranes). After the differentially permeabilized astrocytes had been incubated with the β1-AR antibodies, astrocytes were permeabilized again, this time with triton, to open the nuclear membranes prior to incubation with Lamin A/C antibodies. This allowed identification of nuclei in all astrocytes. For both antibodies, the dense nuclear β1-AR-like immunofluorescence observed in triton-permeabilized astrocytes was almost completely absent in digitonin-permeabilized astrocytes (**Fig. 3C-F, G-J**). To further confirm nuclear localization, we purified plasma membrane and nuclear proteins from primary astrocyte cultures (see Methods) and conducted western blots. Immunoreactivity for the nuclear lamins A and C appeared in whole cell lysates and nuclear fractions, but not in plasma membrane fractions, while immunoreactivity for the ATP1A1 subunit of the sodium-potassium ATPase appeared in whole cell lysates and plasma membrane fractions, but not in nuclear fractions (**Fig. 4A**). β1-AR-like-immunoreactive bands of approximately 65 kDa molecular weight, consistent with previous studies (Clements and Jamali, 2009; Hakalahti et al., 2010) were observed in whole cell lysates, as well as in both plasma membrane and nuclear fractions (**Fig. 4B**).

**Fig. 3.**
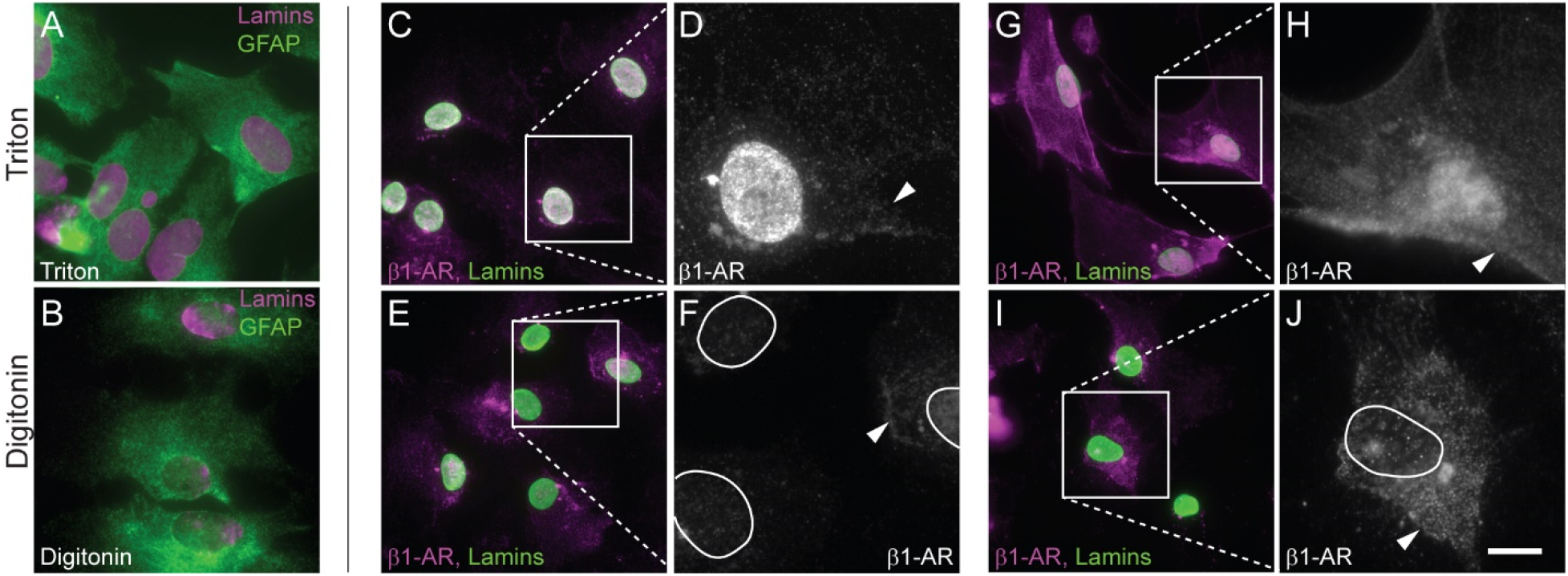
β1-adrenergic receptors are localized to the astrocyte inner nuclear membrane. **(A, B)** Fluorescence photomicrographs of formaldehyde-fixed mouse primary astrocytes permeabilized with Triton X-100 (**A**) or digitonin (**B**) and incubated with antibodies against GFAP (green) and lamin A/C (magenta). (**C-J**) Fluorescence photomicrographs depicting β1-AR (magenta in **C, E, G, I**; white in **D, F, H, J**) and lamin A/C (green in **C, E, G, I**) immunoreactivity in mouse primary astrocytes. Astrocytes were *initially* permeabilized with either Triton X-100 (**C**, **D, G, H**) or digitonin (**E**, **F, I, J**) and incubated with antibodies directed against the extracellular loop 2 (**C-F**) or C-terminal domain (**G-J**) of β1-AR. After β1-AR antibody incubation, all cells were re-permeabilized with Triton X-100 and incubated with an antibody against Lamin A/C (green in **C, E, G,** and **I**) to label nuclei. Arrowheads indicate β1-AR-immunoreactive perikarya. Nuclei are outlined in white in **F** and **J**. Images are representative of n ≥ 3 independent experiments. Scale bar, 25 μm (**A-C, E, G, I**); 9.25 μm (**D**, **F, H, J**).

**Fig. 4.**
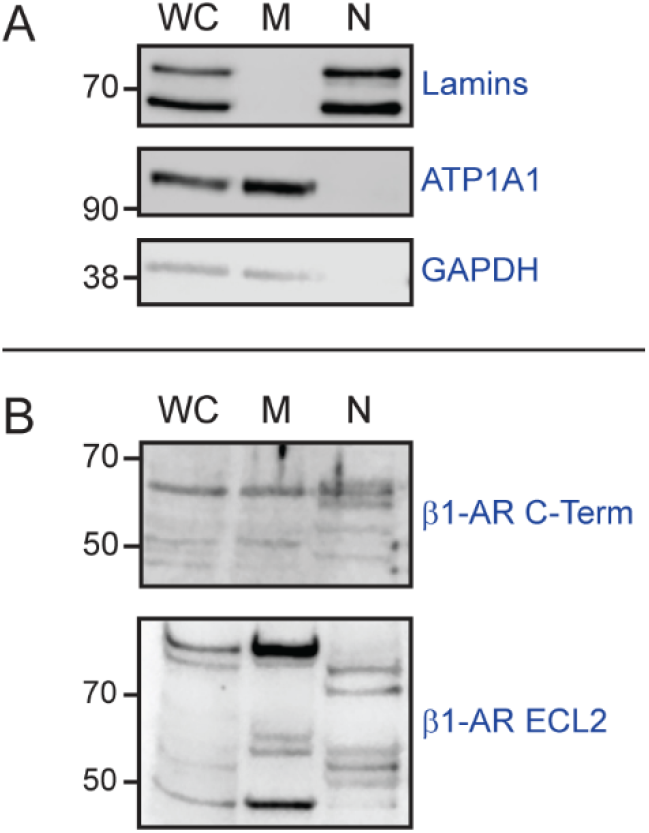
Nuclear localization of β1-adrenergic receptors in astrocytes. (**A**) Validation of subcellular fractionation protocol. Immunoblotting of proteins from whole cell lysate, membrane, and nuclear fractions using antibodies against lamin A/C (Lamins), sodium/potassium ATPase subunit alpha-1 (ATP1A1), and glyceraldehyde 3-phosphate dehydrogenase (GAPDH). Images are representative of n = 4 independent experiments. (**B**) β1-AR immunoblotting of whole cell lysate, membrane, and nuclear proteins from primary cultured astrocytes using antibodies directed against either the C-terminal tail or ECL2 of β1-AR. All images are representative of n = 3 independent experiments.

### G_sα_ and G_s_-coupled signaling components are localized to the nucleus in cultured astrocytes

To begin testing the hypothesis that nuclear membrane-localized β1-ARs are functionally coupled to intranuclear G-protein-mediated signaling pathways, we used immunofluorescence with differential detergent permeabilization to examine the potential nuclear localization of several components of canonical G-protein-coupled receptor signaling. Specifically, we used antibodies against G_αs_, adenylyl cyclase (ADCY) 2, ADCY 5/6, and PKA regulatory subunit isoforms I and II (PKA-RI, PKA-RIIα). Nuclei of cultured astrocytes were identified by immunolabeling with an antibody against nuclear lamins A and C. Decisions of which isoforms of adenylyl cyclase and PKA subunits to examine were based on astrocyte gene expression data from the brain RNASeq database (Zhang et al., 2014).

We observed dense G_sα_-like immunofluorescence over nuclei, and more diffuse, punctate immunofluorescence over somata of triton-permeabilized astrocytes (**Fig. 5A, B**). Nuclear G_sα_-like immunofluorescence was almost completely absent in digitonin-permeabilized astrocytes, while immunofluorescence over the soma remained (**Fig. 5C, D**). In western blots of proteins from astrocyte subcellular fractions, a G_sα_-ir band of approximately 45 kDa was observed in both membrane and nuclear fractions. A 50-kDa G_sα_-ir band was observed in nuclear, but not membrane fractions, and an intermediate sized band was observed in membrane, but not nuclear, fractions (**Fig. 5U**). These bands may represent distinct G_αs_ isoforms that have been previously reported (Robishaw et al., 1986).

**Fig. 5.**
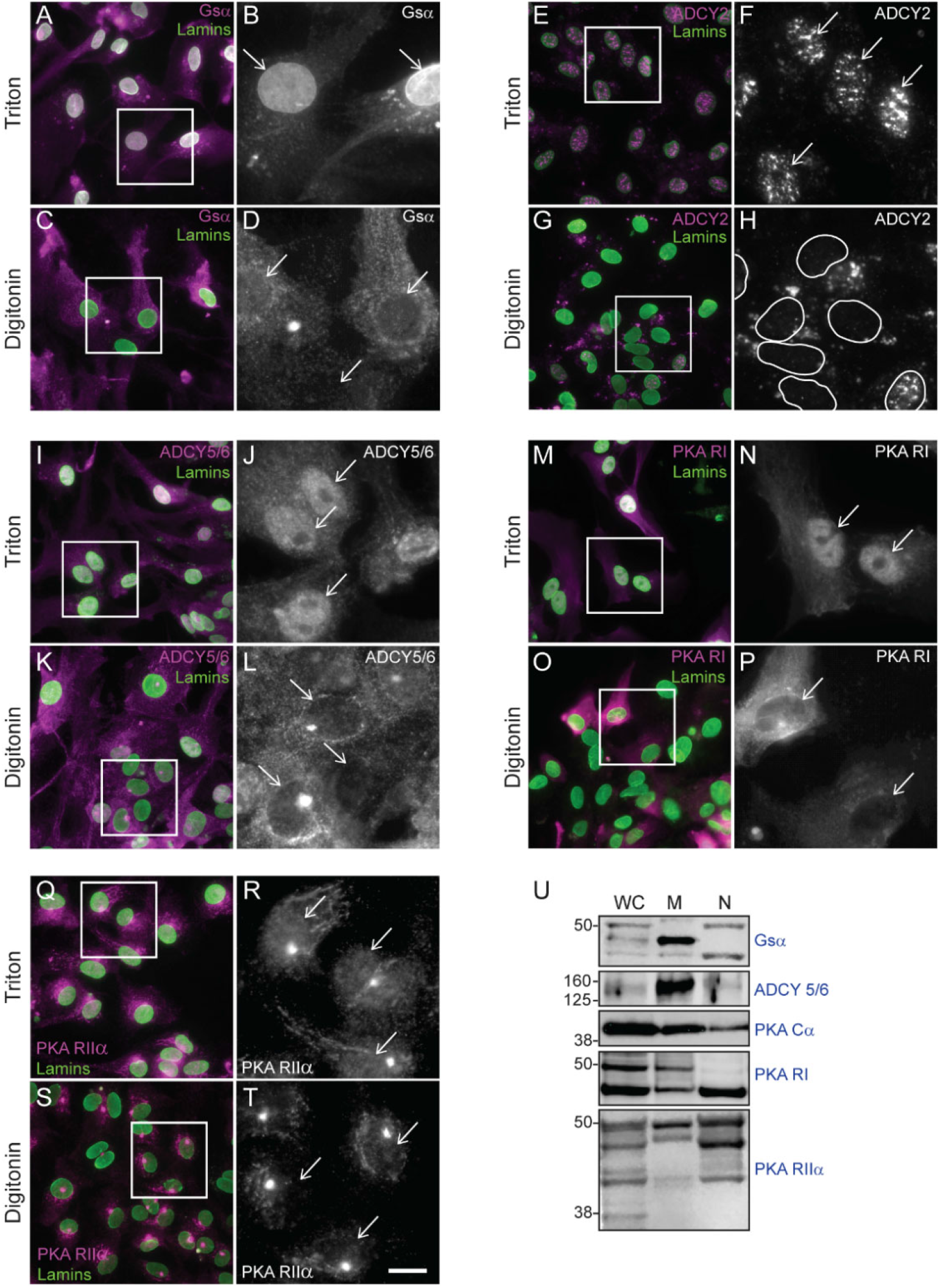
Nuclear localization of β-adrenergic signaling components in mouse astrocytes. (**A**-**T**) Fluorescence photomicrographs of mouse primary astrocytes. Astrocytes were permeabilized with either Triton X-100 or digitonin, followed by incubation with antibodies directed against adrenergic receptor downstream signaling components (magenta in dual color images, white in single color). All astrocytes were then permeabilized with Triton X-100 and incubated with an antibody against lamins A and C (green) to label nuclei. Boxes indicate areas shown at higher magnification to the right. Arrows indicate the position of nuclei. Images are representative of n ≥ 3 independent experiments for each of the following proteins: (**A-D**) Alpha subunit of the stimulatory G protein (G_sα_); (**E-H**) adenylyl cyclase 2 (ADCY2); (**I-L**) adenylyl cyclase 5/6 (ADCY 5/6); (**M-P**) PKA regulatory subunit I (PKA RI); and (**Q-T**) PKA regulatory subunit IIα (PKA RIIα). In each pair of images, scale = 25 μm in the dual-color image, 9.25 μm in grayscale image. (**U**) Immunoblotting of whole cell lysate, membrane, and nuclear proteins from primary mouse astrocytes using antibodies against G_sα_, ADCY 5/6, PKA catalytic subunit (PKA Cα), PKA RI, and PKA RIIα. Images are representative of n = 3 independent experiments.

ADCY2-like immunofluorescence was observed primarily as bright speckles over the nuclei of triton-permeabilized astrocytes, with very little signal observed over somata (**Fig. 5E, F**). Nuclear staining was not observed in digitonin-permeabilized astrocytes (**Fig. 5G, H**). Punctate ADCY5/6-like immunofluorescence was observed diffusely over cell bodies, and densely over nuclei, of triton-permeabilized astrocytes (**Fig. 5I, J**). In digitonin-permeabilized astrocytes, cell body ADCY5/6 immunostaining was observed over cell bodies, but not over nuclei (**Fig. 5K, L**). In western blots of astrocyte subcellular fractions, ADCY 5/6-ir bands with apparent molecular weight between 125 and 160 kDa were observed in whole cell, plasma membrane, and nuclear fractions (**Fig. 5U**). (Predicted molecular weights = 130 kDa (ADCY6 UniProt ID: Q01341), 139 and 148 kDa (ACDY5 UniProt ID: P84309)).

PKA-RI-like immunoreactivity was observed over cell bodies and nuclei of triton-permeabilized astrocytes, with dense staining observed over nuclei (**Fig. 5M, N**). Nuclear PKA-RI immunostaining was almost completely absent in digitonin-permeabilized astrocytes, while cell body staining remained (**Fig. 5O, P**). In western blots, a PKA-RI-ir band of approximately 40 kDa was observed in both membrane and nuclear fractions, while a 50-kDa band was observed only in membrane fractions (predicted molecular weight = 43 kDa UniProt ID: Q9DBC7) (**Fig. 5U**). Punctate PKA-RIIα-like immunoreactivity was distributed evenly over cell bodies and nuclei of triton-permeabilized astrocytes, and at higher density in endomembrane (ER or Golgi)-like profiles (**Fig. 5Q, R**). Nuclear PKA-RIIα immunostaining was weaker in digitonin-permeabilized than in triton-permeabilized astrocytes (**Fig. 5S, T**). In western blots, PKA-RIIα-ir bands of 40-45 kDa (predicted molecular weight = 45 kDa, UniProt ID: P12367) were observed in whole cell lysate, membrane, and nuclear fractions (**Fig. 5U**).

We observed PKA catalytic subunit (alpha isoform) immunoreactivity in western blots of astrocyte nuclear and membrane fractions. A PKA Cα-immunoreactive band of approximately 40 kDa molecular weight (predicted molecular weight = 40 kDa, UniProt ID: P05132) was observed in whole cell lysates, plasma membrane, and nuclear fractions (**Fig. 5U**).

### Catecholamine transporters are localized to astrocyte nuclei and plasma membranes

Access of norepinephrine, which is cationic and membrane-impermeable at physiological pH, to β1-ARs localized to inner nuclear membranes would require carrier-mediated transport across both plasma and outer nuclear membranes. While our studies indicate that OCT3 could potentially mediate transport across both membranes, it is possible that additional catecholamine transporters are involved. To characterize more completely the mechanisms by which norepinephrine may access nuclear β1-AR, we examined the localization of two additional uptake_2_ transporters: organic cation transporter 2 (OCT2) and the plasma membrane monoamine transporter (PMAT), and of the uptake_1_ catecholamine transporters NET (norepinephrine transporter) and DAT (dopamine transporter), using immunofluorescence (exclusively on triton-permeabilized astrocytes) and western blot.

Both OCT2-like and PMAT-like-ir were observed over nuclei and somata of triton-permeabilized astrocytes (**Fig. 6A-D**). In western blots, OCT2-ir bands of between 50 and 70 kDa molecular weight (predicted molecular weight = 62 kDa (Karbach et al., 2000; Urakami et al., 1999)) were observed in membrane and nuclear fractions (**Fig. 6I**). PMAT-ir bands of approximately 50 and 60 kDa molecular weight (predicted molecular weight = 58 kDa (Xia et al., 2007)) were observed in nuclear fractions, while a single ~65-kDa PMAT-ir band was observed in membrane fractions (**Fig. 6I**). Faint NET-like immunofluorescence was observed over astrocyte somata and nuclei (**Fig. 6E, F**). In western blots, NET-like-ir bands of approximately 45-, 65- and 70-kDa molecular weight were observed in both membrane, and nuclear fractions, though all bands in membrane fractions were very faint (**Fig. 6I**). Non-glycosylated hNET migrates in western blots at approximately 46-50 kDa (Melikian et al., 1996), with glycosylated forms appearing at 54- and 70 kDa (Matthies et al., 2009; Melikian et al., 1996). Faint DAT-like immunofluorescence was observed over astrocyte cell bodies and nuclei (**Fig. 6G, H**). A major 50-kDa DAT-ir band was observed in western blots of astrocyte nuclear fractions, with additional bands of higher molecular weight in both nuclear and membrane fractions (**Fig. 6I**). Fully glycosylated DAT appears in western blots as a broad band centered around 75-85 kDa molecular weight, while non-glycosylated DAT (40-50 kDa), and several intermediate glycosylation states of DAT (40-85 kDa) are also observed (Li et al., 2004).

**Fig. 6.**
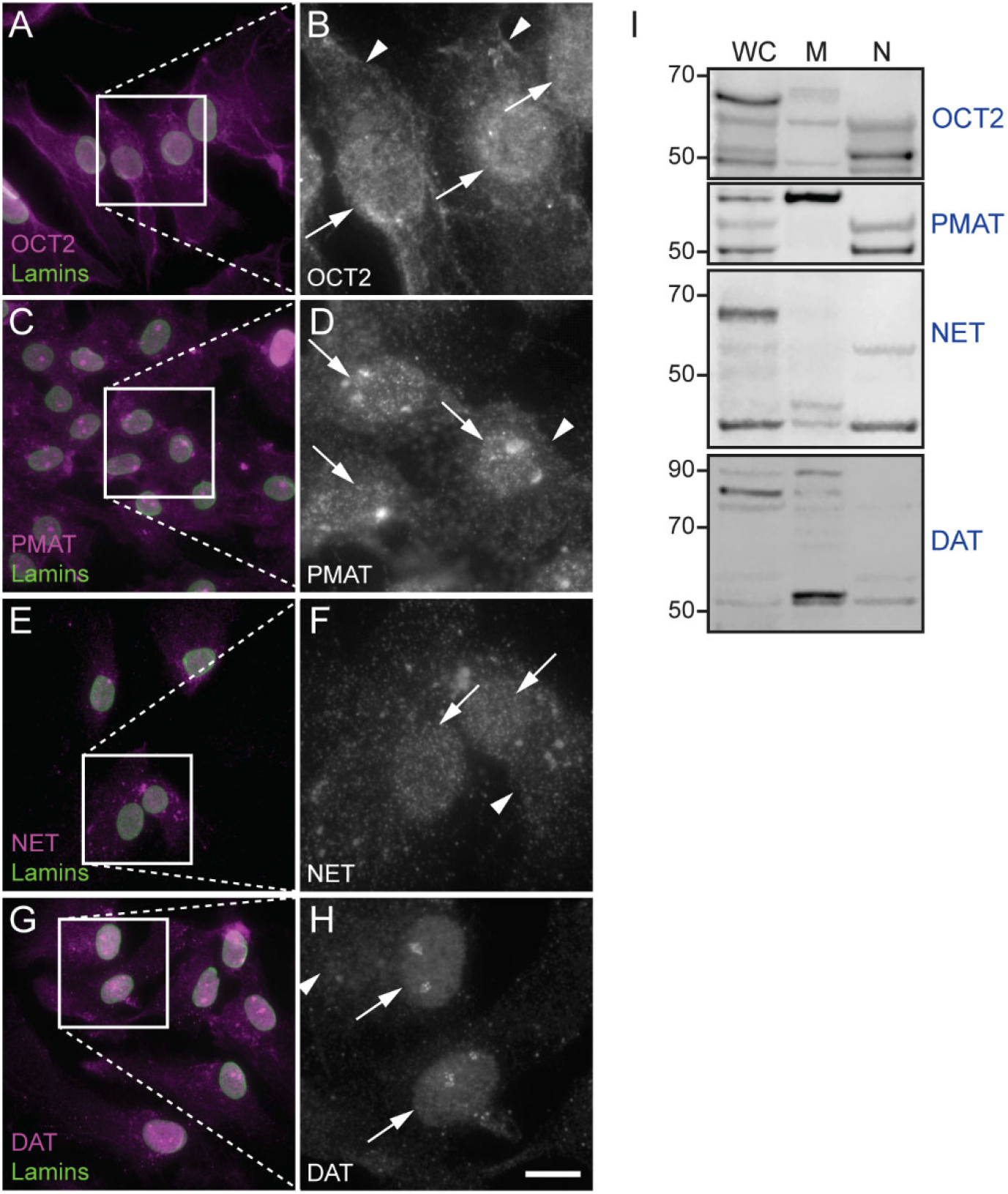
Localization of catecholamine transporters in mouse astrocytes. (**A**-**H**) Fluorescence photomicrographs of mouse primary astrocytes permeabilized with Triton X-100 and incubated with antibodies against lamin A/C (green) and catecholamine transporters (magenta and white). Images are representative of n ≥ 3 independent experiments for each of the following transporters: (**A, B**) organic cation transporter 2 (OCT2); (**C, D**) plasma membrane monoamine transporter (PMAT); (**E, F**) norepinephrine transporter (NET); (**G, H**) dopamine transporter (DAT). Arrows indicate the positions of nuclei. Arrowheads indicate cell body immunoreactivity. Scale bar, 25 μm in all dual-color images, 9.25 μm in all grayscale images. (**I**) Immunoblotting of whole cell lysate, membrane, and nuclear proteins from primary mouse astrocytes using antibodies against OCT2, PMAT, NET and DAT. Images are representative of n = 3 independent experiments.

### Norepinephrine induces rapid and prolonged increases in nuclear PKA activity

The localization of β1-ARs to inner nuclear membranes and the evidence that G-proteins and their signaling partners occur in astrocyte nuclei suggest that nuclear receptors are capable of activating downstream cAMP signaling. The presence of OCT3 and other catecholamine transporters in both plasma and nuclear membranes suggests the hypothesis that access to and activation of nuclear membrane β1-ARs is gated by transporter activity. To determine whether nuclear β1-ARs are functionally coupled to signaling machinery and to test the hypothesis that activation of nuclear receptors is transporter-gated, we transfected astrocytes with plasmids driving the expression of a nuclear localized PKA activity sensor, ExRai-AKAR-NLS, to allow monitoring of nuclear PKA activity in real time. This sensor consists of a PKA substrate sequence and a phosphoamino acid-binding domain fused with the circularly permuted GFP from GCaMP3. This recombinant protein displays two excitation peaks (~400 and ~509 nm with a shoulder at ~480 nm), both of which emit at ~515 nm. Phosphorylation of the PKA substrate sequence induces a conformational change in the sensor that results in a decrease in 400 nm-induced emission and an increase in 488 nm-induced emission, resulting in a large increase in the “excitation ratio” (Mehta et al., 2018). We examined the effects of bath-applied norepinephrine (50 nM) on nuclear excitation ratios in astrocytes in the presence or absence of catecholamine transport inhibitors. Transfected astrocytes were pretreated with either vehicle or a cocktail of catecholamine transport inhibitors including atomoxetine (NET inhibitor), GBR12909 (DAT inhibitor), and corticosterone at a concentration shown to inhibit OCT3, OCT2, and PMAT (Duan and Wang, 2010; Engel et al., 2004; Gründemann et al., 1998). After the preincubation period, excitation ratiometric data were collected every 20 seconds for a 3-minute baseline period, after which norepinephrine was bath applied to all astrocytes and ratiometric data were collected every 20 seconds for an additional 30 minutes. Increases in nuclear PKA were detected as increases in the excitation ratio. Norepinephrine-induced increases in nuclear excitation ratio were observed in both vehicle- and transport inhibitor-pretreated cells, but the kinetics of responses in the two groups were different. In vehicle-pretreated cells, NE treatment produced rapid and robust increases in nuclear excitation ratio over the initial 10-minute period (representative traces and ratio images in **Fig. 7A, B**; mean responses in **Fig. 7C, D**). In transport inhibitor-treated cells, NE treatment led to delayed increases in nuclear excitation ratio that appeared in the second 10-minute post-treatment period. A 2 (Group: Vehicle vs Inhibitors) x 4 (Period: Baseline vs 0-10 vs 10-20 vs 20-30 minutes) ANOVA revealed higher average ratios in the Vehicle group relative to the Inhibitors group, *F*(1, 34) = 8.18, *p* = 0.007, and an increase in the ratio across time periods, *F*(1, 102) = 64.93, *p* < 0.0001, as well as an interaction between group and time period, *F*(1, 102) = 3.31, *p* = 0.023 (**Fig. 7E**). Planned interaction contrasts showed that excitation ratios increased more in the Vehicle group from baseline to 0-10 minutes relative to the Inhibitors group, *F*(1, 34) = 38.47, *p* < 0.0001, consistent with the hypothesis that the rapid nuclear PKA response is dependent on catecholamine transport. However, across the subsequent time periods (0-10 minutes vs. 10-20 minutes), the rate of change in nuclear PKA was similar between the two groups (*F*(1, 34) = 0.89, *p* = 0.35). A Tukey’s HSD analysis supported this conclusion, showing that excitation ratios increased across every post-NE time period in the vehicle-pretreated group (*p*s < 0.025), but only exhibited a significant elevation above baseline after 10-minutes of norepinephrine exposure in the transport inhibitor-pretreated group (*p*s < 0.004). Taken together, the results suggest that rapid nuclear PKA responses (within 10 minutes) in astrocytes are dependent on carrier-mediated transport of norepinephrine, and that more gradual increases in nuclear PKA signaling (after 10 minutes) are not.

**Fig. 7.**
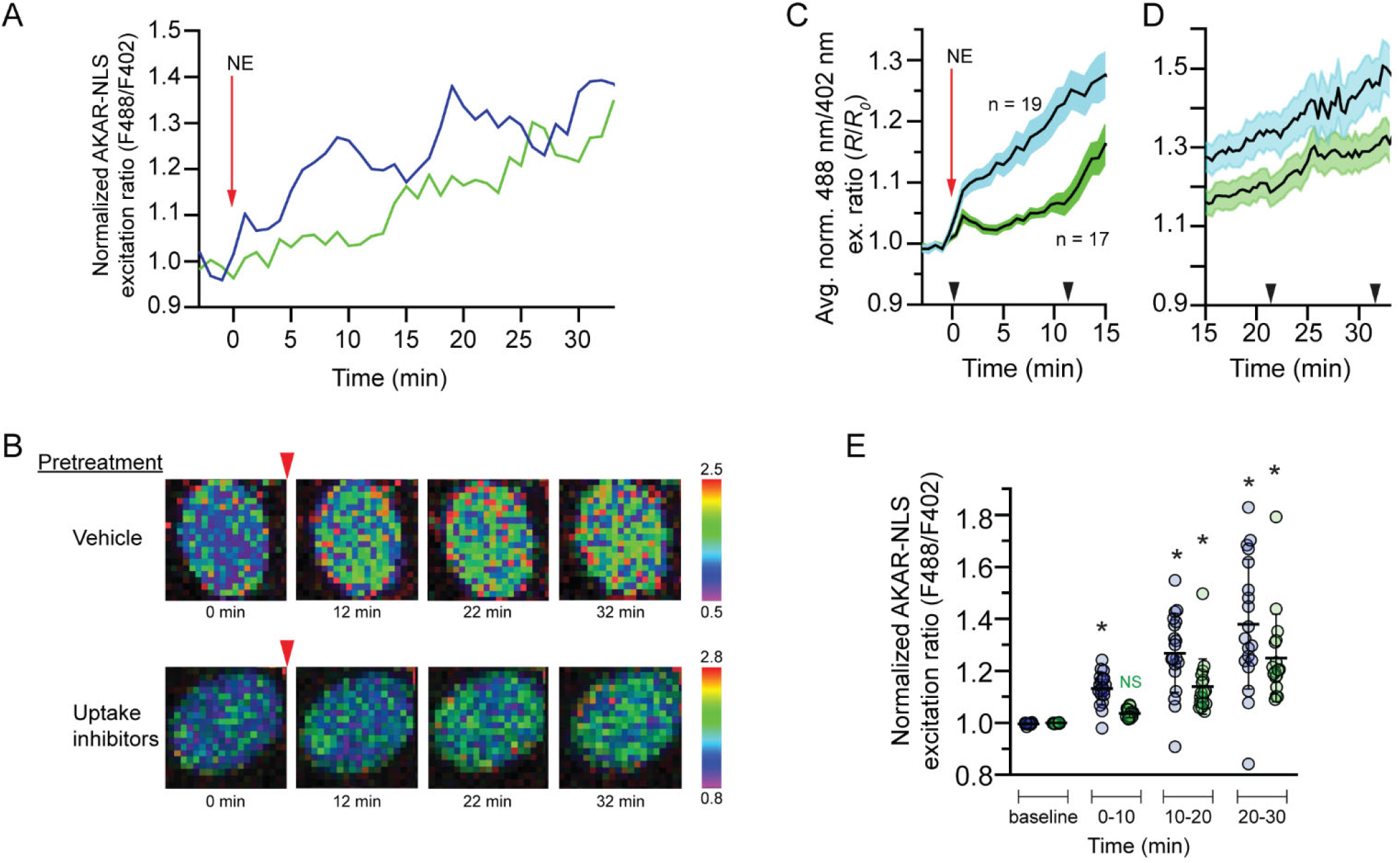
Rapid norepinephrine-induced increases in nuclear PKA activity require catecholamine transporter function. (**A**) Representative time courses of nuclear PKA responses to 50 nM norepinephrine in ExRai-AKAR-NLS-expressing astrocytes pretreated with vehicle (blue) or catecholamine transport inhibitors (green). Curves are plotted as 488/402 nm excitation ratios normalized with respect to time 0. (**B**) Pseudocolor images of the ExRai-AKAR-NLS ratio responses of the astrocytes from **A**. Warmer colors indicate higher excitation ratios as shown in the scales to the right of the images. Red arrowheads indicate the time of norepinephrine application. Pretreatment condition is indicated to the left of the images. (**C, D**) Average time courses (t = −3 to 15 min in **C**; 15-35 min in **D**) of nuclear PKA responses to 50 nM norepinephrine in ExRai-AKAR-NLS-expressing astrocytes pretreated with vehicle (blue, n = 19 cells from 5 separate experiments) or transport inhibitors (green, n = 17 cells from 5 separate experiments). Curves are plotted as 488/402 nm excitation ratios normalized with respect to time 0. Solid lines represent the mean; shaded areas represent s.e.m. (**E**) Normalized nuclear PKA responses to 50 nM norepinephrine of vehicle (blue)- or transport inhibitor (green)-pretreated ExRai-AKAR-NLS-expressing astrocytes (same cells shown in (**C**)). Ratiometric responses obtained during the indicated time periods were pooled and averaged for each cell. Thick, horizontal lines represent the mean and error bars represent SD. * indicates significantly different from the baseline excitation ratio for that group (*p* < 0.05).

## DISCUSSION

Norepinephrine is a key mediator of integrated central nervous system responses to stressful and arousing stimuli. In addition to its well-characterized influences on neuronal function and synaptic plasticity, norepinephrine has powerful effects on core astrocyte functions, including neuroprotective (Junker et al., 2002), immunomodulatory (Laureys et al., 2014), and metabolic support functions (Coggan et al., 2018). These effects, mediated by G-protein-coupled α- and β-adrenergic receptors, include both rapid, short-term changes in cellular physiology and delayed, long-term changes in gene expression and cellular structure. For example, norepinephrine induces rapid activation of cytosolic glycogen phosphorylase, leading to increases in glycogen breakdown and lactate release (Coggan et al., 2018; Hertz et al., 2010; Sorg and Magistretti, 1991) and gradual increases in the expression of glycogen synthase mRNA and protein, allowing adaptive re-synthesis of glycogen (Sorg and Magistretti, 1991). Thus, integrated cellular responses to norepinephrine require the propagation of adrenergic receptor signaling events to cytosolic, plasma membrane, and nuclear targets. Dysregulation of noradrenergic signaling and of astrocyte function have been implicated in neurodegenerative disorders, including Alzheimer’s disease and multiple sclerosis (Keyser et al., 2010; Santello et al., 2019). Determining the mechanisms by which norepinephrine-induced signals reach distinct cellular locations is critical for a complete understanding of noradrenergic regulation of cellular function under normal and pathological conditions.

The present studies provide evidence that the inner nuclear membrane is an adrenergic signaling platform in astrocytes. They reveal a population of functional, G-protein-coupled β1-adrenergic receptors localized to the inner nuclear membranes of astrocytes and indicate that norepinephrine accesses these receptors through the actions of catecholamine transporters localized to plasma and outer nuclear membranes. Norepinephrine-induced activation of these receptors leads to rapid increases in nuclear PKA activity that are blocked by catecholamine uptake inhibitors. These receptors represent a powerful mechanism by which NE may directly influence nuclear processes including transcription factor activity and chromatin modification, leading to profound effects on gene expression, and contributing to the neuroprotective, immunomodulatory, and metabolic regulatory actions of norepinephrine in astrocytes.

Growing numbers of studies have demonstrated that adrenergic and other G-protein-coupled receptors can initiate signaling cascades from intracellular locations including the nuclear membrane (Campden et al., 2015; Jong et al., 2018). Our studies provide evidence, using western blot on subcellular fractions as well as immunofluorescence, that β1-ARs are localized to both plasma and nuclear membranes in astrocytes. They further indicate that β1-ARs are localized to inner, but not outer, nuclear membranes of cultured astrocytes, as the nuclear immunofluorescence observed in triton-permeabilized astrocytes with antibodies directed against both intra- and extracellular epitopes of the receptor was attenuated when astrocytes were permeabilized with digitonin. If nuclear labeling represented receptors localized to the outer nuclear membrane or to peri-nuclear ER, nuclear fluorescence resulting from at least one of the two antibodies would be unchanged by digitonin permeabilization. These findings are the first to document nuclear localization of adrenergic receptors in any CNS cell type and suggest that, in addition to its actions at the plasma membrane, norepinephrine may initiate G_s_-mediated signaling cascades in the nuclear compartment of astrocytes. They are consistent with studies demonstrating that, in cardiomyocytes, α- and β- adrenergic receptors are localized to the nuclear envelope, where they activate G_q_- and G_s_-coupled signaling pathways and mediate effects on gene expression and myocyte physiology (Vaniotis et al., 2011; Wu et al., 2014).

Our studies suggest that, while β1-ARs are localized to both cell surface and nuclear membranes, receptors at the two sites may initiate distinct cAMP signaling processes. The subcellular localizations of key components of the canonical G_s_ signaling machinery, including G_αs_, adenylyl cyclase isoforms, and PKA subunits, differed between nuclei and cell bodies. Distinct isoforms of adenylyl cyclase and PKA regulatory subunits were observed in nuclear and plasma membrane sites, consistent with the emerging understanding that PKA can be organized into signaling microdomains organized by A-Kinase-Anchoring Proteins (AKAPs) with distinct cellular localizations (Torres-Quesada et al., 2017). Thus, nuclear and plasma membrane-localized receptors may initiate distinct cAMP signaling processes which mediate distinct aspects of the overall response to norepinephrine.

The immunostaining patterns observed in the present studies suggest that there are robust, and possibly diverse, cAMP signaling mechanisms in astrocyte nuclei. G_αs_ and at least two distinct isoforms of adenylyl cyclase were identified in astrocyte nuclei, suggesting that activation of nuclear β1-ARs may induce rapid increases in nuclear cAMP as has been previously described in HEK-293 cells (Sample et al., 2012). The western blot banding patterns indicate that there may be distinct G_αs_ isoforms localized to nuclear and plasma membrane sites. At least two, and possibly three, isoforms of adenylyl cyclase were identified in astrocyte nuclei. ADCY2 was observed exclusively in nuclei, where it was localized to discrete, brightly labeled clusters, while ADCY5 and/or 6 were localized to both plasma and nuclear membranes. The ADCY2-labeled nuclear structures may represent “speckles”, nuclear microdomains enriched in RNA processing enzymes, RNA Polymerase II, and certain transcription factors (Spector and Lamond, 2011), and recently shown to be enriched with AKAP95 and other signaling proteins (Li et al., 2020). It was not possible to determine from our data whether staining in either subcellular compartment represented adenylate cyclase 5, 6 or a combination of the two isoforms. The finding of nuclear-localized adenylyl cyclase is consistent with early studies that identified, in purified lymphocyte nuclei, adenylyl cyclase activity that could be stimulated by treatment with the β-AR agonist isoproterenol (Wedner and Parker, 1977), suggesting that lymphocytes may also express nuclear-localized β-ARs.

Consistent with previous studies that demonstrated the existence of a cAMP-activable pool of PKA holoenzyme resident in nuclei of HEK-293 cells (Clister et al., 2019; Sample et al., 2012), our studies suggest that a subset of PKA holoenzyme resides in the nuclei of unstimulated astrocytes. Both PKA RI and RIIα were localized to astrocyte nuclei, with RI more strongly localized to nuclei and RIIα distributed more evenly between plasma membrane and nuclei. PKA RIIα also appeared in putative endomembrane profiles, and was densely localized to a perinuclear structure, likely the centrosome. PKA RIIα has previously been localized to both centrosomes and Golgi apparatus (Keryer et al., 1999).

The present studies provide evidence that multiple catecholamine transporters may gate access of norepinephrine to nuclear receptors. They confirm our previous observation that OCT3 is localized to plasma and outer nuclear membranes (Gasser et al., 2017), and they suggest potential roles for OCT2 and PMAT, as well as the uptake_1_ transporters DAT and NET, in gating activation of nuclear receptors. They are consistent with studies demonstrating that inhibition or genetic knockout of OCT3 decreases the activation of nuclear alpha-adrenergic receptors (Wright et al., 2008) and Golgi-localized β1-ARs (Irannejad et al., 2017; Wang et al., 2020). While the present studies are the first to suggest nuclear membrane localization of OCT2, PMAT, and NET, they are not the first to suggest that DAT may be localized to the nucleus. Immunogold electron microscopic examination of neurons in the substantia nigra revealed DAT localized to both inner and outer nuclear membranes (Block et al., 2015; Hersch et al., 1997). Our studies did not determine the specific localization of any transporters to inner or outer nuclear membranes and provide no information about the orientation of any of the transporters in the membrane. The role of a given transporter in either allowing or limiting the activation of nuclear β1-ARs would depend on its localization to the inner or outer nuclear membrane and on its orientation in those membranes. Localization of monoamine transporters to the nuclear envelope is also interesting in light of recent studies documenting the phenomenon of histone monoaminylation, in which dopamine and serotonin, both substrates of the uptake_2_ transporters, are enzymatically conjugated to histones and modulate gene expression (Chan and Maze, 2020; Farrelly et al., 2019).

We observed two classes of nuclear PKA responses to norepinephrine based on their sensitivity to pretreatment with inhibitors of catecholamine transport: a transporter-dependent, early response, and a transporter-independent, delayed response. The transporter-dependent response was rapid, detectable within the first minutes of norepinephrine treatment, and was absent in cells pre-treated with uptake inhibitors. The transporter-independent response was more gradual, only becoming significant during the second 10-minute time block after norepinephrine treatment and was unaffected by uptake inhibition. The rapid onset of early, transporter-dependent responses suggests that they are mediated by PKA that is resident to the nucleus and activated in response to nuclear β1-AR-initiated G_s_ signaling, and not by PKA activated at the plasma membrane and diffusing to the nucleus. This is consistent with studies examining the kinetics of nuclear PKA activation in response to cAMP generated either at the plasma membrane or in the nucleus (Sample et al., 2012). In those studies, when cAMP was generated at the plasma membrane, nuclear PKA activity increased gradually, with a half time (t_1/2_) of approximately 20 minutes and was mediated by catalytic subunits which had been activated at the plasma membrane and diffused into the nucleus. When cAMP was generated within the nucleus, nuclear PKA activity increased rapidly, with t_1/2_ of 3-4 minutes, and appeared to be mediated by nuclear-resident PKA. In our studies, the rapid onset of the transporter-dependent increase in PKA activity suggests that it reflects activation of nuclear-resident PKA in response to locally generated cAMP. The delayed onset of the transporter-independent PKA response in the present studies suggests that it reflects the activity of cytosolic PKA activated in response to plasma-membrane-localized β-AR receptor activation. Alternatively, it is possible that the inhibition of norepinephrine uptake in our experiments facilitated the activation of cell surface α2-adrenergic receptors which have been shown to inhibit β-AR-induced activation of adenylyl cyclase (Northam et al., 1989). This could explain the delayed increase in nuclear PKA activity we observed. Further studies are needed to explore this possibility.

Norepinephrine exerts powerful and pervasive actions in the central nervous system. In astrocytes, these effects include neuroprotective, immunomodulatory, and metabolic regulatory actions. Many of these actions critically involve regulation of nuclear processes and alterations in gene expression patterns. Activation of astrocyte β-adrenergic receptors induces phosphorylation of RNA Polymerase II (Lee and Jungmann, 1981), histones (Harrison et al., 1980), and CREB (Koppel et al., 2018), and regulates the expression of neurotrophic factors (Juric et al., 2008; Koppel et al., 2018), metabolic enzymes (Allaman et al., 2000; Pellegri et al., 1996) and immunoregulatory proteins (Feinstein et al., 2002; Gavrilyuk et al., 2002; Madrigal et al., 2009). Our findings suggest a novel and powerful mechanism by which norepinephrine may initiate these and other actions in the nucleus. They add to the growing body of evidence that G-protein-coupled receptors can be localized to, and activated at, endomembrane sites (Jong et al., 2018). The presence of β1-ARs at both nuclear and plasma membranes suggests the possibility that the two populations of receptor may mediate distinct aspects of the integrated cellular response to norepinephrine, each contributing uniquely to the neuroprotective, and immunomodulatory and metabolic regulatory actions of norepinephrine in astrocytes.

## MATERIALS AND METHODS

### Antibodies and plasmids

Antibodies used in western blot and immunofluorescence studies are described in **Table 1**. The PMAT antibody was the generous gift of Dr. Joanne Wang. The pcDNA3 Flag β1-adrenergic-receptor plasmid was a gift from Dr. Robert Lefkowitz (Addgene plasmid # 14698; http://n2t.net/addgene:14698 ; RRID:Addgene_14698). The ExRai-AKAR-NLS plasmid was a gift from Dr. Jin Zhang.

**Table 1.**
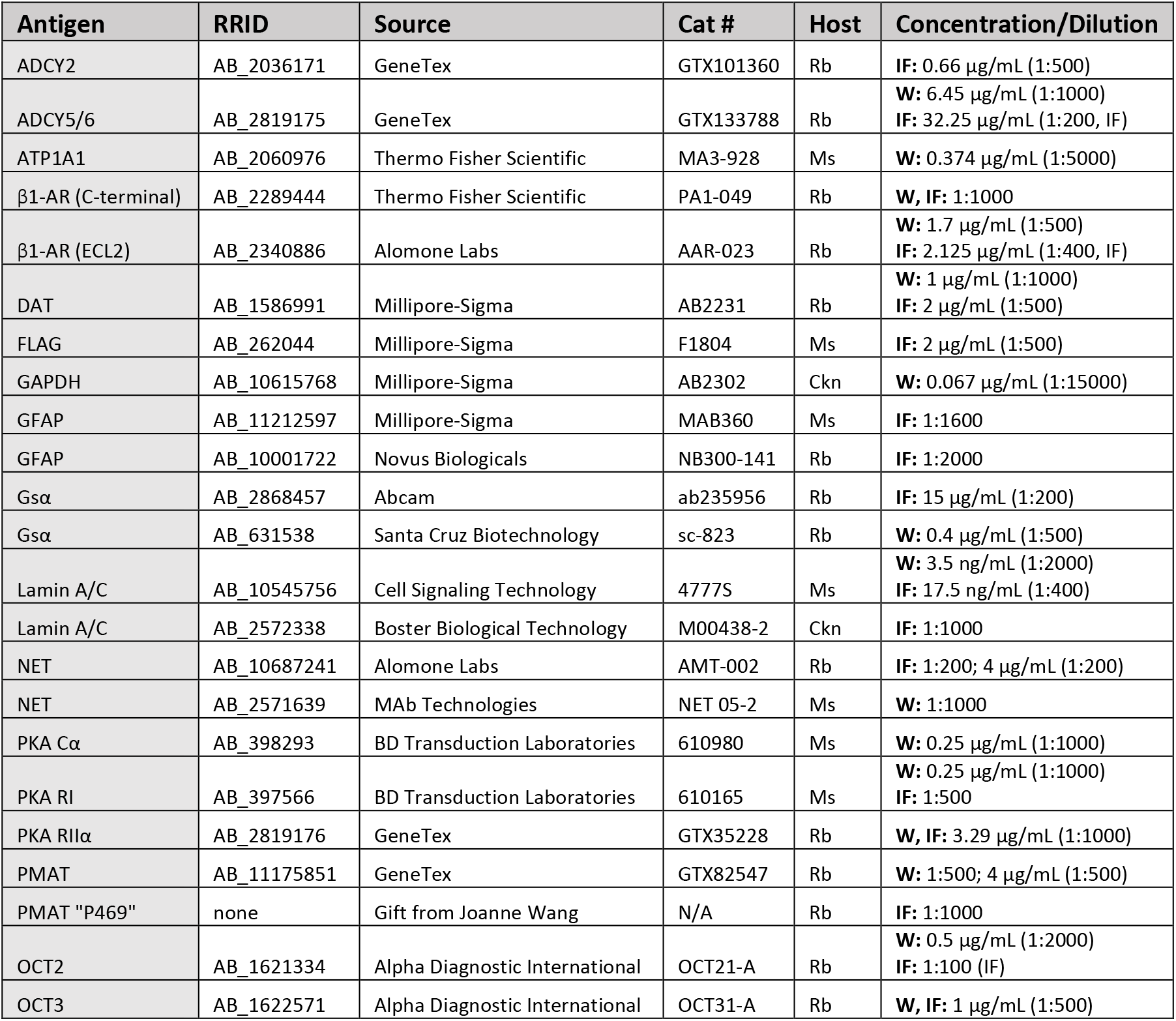
Primary antibodies used in western blot and immunofluorescence experiments. W: Western blot; IF: Immunofluorescence.

### Primary astrocyte cell culture

Astrocytes were cultured from cerebral cortices of postnatal day 1-2 C57BL/6 mouse pups bred in an in-house colony. Mice were handled in accordance with a protocol approved by the Marquette University institutional animal care and use committee and in compliance with the US National Institutes of Health Guide for the Care and Use of Laboratory Animals. Primary cortical astrocyte cultures and reagents were prepared as described in Uliasz et al (Uliasz et al., 2011). After 10 days in culture, confluent cells were treated with an anti-mitotic agent, cytosine arabinoside (ara-C), to limit microglial contamination. Astrocytes were harvested for subcellular fractionation or plated for immunofluorescence or excitation ratio imaging 3-5 weeks after initial culture. One day before all experiments astrocytes were treated with L-leucine methyl ester (LME) to further deplete microglia.

### Immunofluorescence and microscopy

For all immunofluorescence experiments, cultured astrocytes were plated on glass-bottomed chamber slides (MatTek, Ashland, MA, USA or Millicell EZ Slide, Millipore Sigma). 5-7 days after plating, astrocytes were fixed (PBS + 2% paraformaldehyde, 10 min, 4°C), rinsed in 0.05M PBS, and permeabilized by incubation in blocking buffer (0.05M PBS, 0.3M glycine, 5% donkey serum) containing one of two detergents: Triton X-100 (0.1%; PBST) or digitonin (10μg/mL, 0.001%). Triton X-100 permeabilizes all cellular membranes, while digitonin, at the selected concentration, preferentially permeabilizes cholesterol-rich membranes, leaving nuclear membranes essentially intact (Jamur & Oliver, 2009). Astrocytes were then rinsed and incubated (overnight, 4°C) in 0.05M PBS containing the primary antibody of interest (see **Table 1** for antibody and dilution information). After thorough rinsing, astrocytes were incubated (2 h, room temp) in 0.1% PBST containing the appropriate fluorophore-conjugated secondary antibody (AlexaFluor 488 or 594, 1:2000; Jackson ImmunoResearch). Triton X-100 was included to allow secondary antibody to reach all cellular compartments. In experiments in which a second antigen was labeled, cells were thoroughly rinsed and incubated in blocking buffer with 0.1% Triton X-100 (20 min, room temperature) prior to incubation with primary and secondary antibodies as above. After final rinses, cells were dried and slides were coverslipped using fluorescent mounting medium (VectaShield, Vector Laboratories, Inc., Burlingame, CA, USA; or EverBrite, Biotium, Inc, Fremont, CA USA). In some studies, nuclei were counterstained using DAPI. Each detergent comparison experiment was repeated at least three times using cells cultured from separate mouse litters. Photomicrographs were captured using a Nikon 80i microscope fitted with an Orca Flash 4.0L T digital camera (Hammamatsu, Japan) linked to a computer running NIS Elements BR software (Nikon Instruments, Melville, NY) or a Nikon A1R+ laser scanning confocal microscope linked to a computer running NIS Elements AR software. Photomicrographs of cells in the two detergent conditions were captured with identical exposure times or, in the case of the confocal microscope, identical laser power and illumination settings. To confirm that the two detergents differentially permeabilized plasma and nuclear membranes, we probed triton- and digitonin-permeabilized astrocytes with anti-lamin A/C antibodies. Dense lamin-like immunoreactivity was observed over the nuclei of triton-permeabilized astrocytes but was nearly absent in digitonin-permeabilized astrocytes (**Fig. 3 A, B**).

#### Localization of FLAG-tagged β1-AR

To confirm the subcellular localization of β1-AR, astrocytes were chemically transfected using Lipofectamine 3000 (Invitrogen) with a plasmid directing expression of a recombinant FLAG-tagged human β1-AR. During transfection cells were incubated with 1 μg of plasmid (37°C, 5 hours), followed by rinsing and incubation for 72 h in fresh growth medium. Cells were then fixed and permeabilized with 0.1% PBST. Immunofluorescence using an anti-FLAG primary antibody (**Table 1**) was carried out as described above.

#### Perfusions and immunofluorescence in mouse brain tissue

Mice (C57BL6, Envigo) were deeply anesthetized by intraperitoneal injection of sodium pentobarbital (100 mg/kg) and were transcardially perfused with ice-cold 0.05 M phosphate-buffered saline followed by 4% paraformaldehyde in 0.1 M sodium phosphate buffer (PB, pH 7.4). Following perfusion, brains were removed and post-fixed in the 4% paraformaldehyde solution for 12 hours at 4°C and rinsed twice in 0.1 M PB for 12 hours. Brains were then incubated in 30% sucrose in 0.1 M PB for approximately 72 hours followed by rapid freezing in dry-ice-chilled liquid isopentane and storage at −80 °C until sectioning. Forebrain sections (25 μm) were cut across the coronal plane using a cryostat (Leica Biosystems, Buffalo Grove, IL, USA), and stored in cryoprotectant (30% ethylene glycol (w/w)/20% glycerol (w/w) in 0.05 M PB, pH 7.4) at −20 °C until immunostaining.

After rinsing in PBS, sections were incubated overnight with C-terminal-directed anti-β1-AR antibody in 0.1% PBST. Sections were rinsed the next day and incubated with fluorophore-conjugated secondary antibodies (AlexaFluor 594-conjugated donkey anti-rabbit IgG; 1:2000; Jackson ImmunoResearch) for 2h. Sections were then rinsed and incubated with anti-GFAP antibody in 0.1% PBST overnight. The following day sections were rinsed, incubated 2h with fluorophore-conjugated secondary antibody (AlexaFluor 488-conjugated donkey anti-mouse IgG), rinsed, and mounted onto SuperFrost microscope slides. After drying, sections were coverslipped with mounting medium (VectaShield + DAPI, Vector Laboratories, Inc., Burlingame, CA, USA). Photomicrographs were acquired using a Nikon 80i microscope fitted with an ORCA-Flash 4.0LT digital camera (Hammamatsu, Japan) linked to a computer running NIS Elements-BR software (Nikon Instruments, Melville, NY).

### Subcellular fractionation, gel electrophoresis and Western blot

Cytosolic, nuclear, and plasma membrane proteins were purified from cultured astrocytes using a commercially available kit (Qproteome Cell Compartment Kit; Qiagen, Inc, Germantown, MD, USA) according to the manufacturer’s instructions. Protein concentration in each fraction was determined (Pierce BCA Protein Assay, ThermoFisher Scientific, Waltham, MA, USA). Subcellular fractions were prepared for electrophoresis by addition of 4x Bolt LDS sample buffer and reducing agent (Invitrogen, Carlsbad, CA, USA) followed by heating at 37°C for 30 min. Proteins (approximately 6 μg/sample) were electrophoresed on 4-12% Bis-Tris polyacrylamide gels (Invitrogen) at 200V for 55 minutes, followed by electroblotting (25V, 0.13A. 17W 1.5 hours) onto Immobilon-FL polyvinylidene difluoride membranes using a wet transfer apparatus (Thermo Fisher). Membranes were dried overnight at room temperature. The following day, membranes were briefly re-wet in 100% methanol, rinsed, blocked (Odyssey TBS Blocking Buffer, Li-Cor Biotechnology, Lincoln, NE), and incubated overnight at 4 °C in blocking buffer containing 0.1% Tween-20 and primary antibodies of interest (see Table S1). Blots were then rinsed and incubated (2 h, room temperature) with species-specific secondary antibodies conjugated to a fluorophore (AlexaFluor 680- or AlexaFluor 800-conjugated goat anti-mouse and/or goat anti-rabbit (Invitrogen); 1:15,000). After rinsing, digital images of the fluorescent bands were captured using an Odyssey Fc Imaging System (LI-COR Biotechnology, Lincoln, NE, USA). Images were captured at each wavelength and saved as separate files. Data for individual antigens are presented separately as black-and-white images. Each immunoblot was repeated at least three times with proteins from separate astrocyte cultures.

### Excitation ratio imaging

Cultured astrocytes were transfected (Lipofectamine 3000; Invitrogen) with a plasmid driving the expression of ExRai-AKAR-NLS, a single-fluorophore biosensor that combines a PKA substrate sequence with the FHA1 domain of AKAR, a circularly permuted GFP, and a nuclear localization sequence. The fluorophore has two discrete excitation peaks: one centered around 400 nm and another around 509 nm, with a shoulder at 480 nm. Phosphorylation of the PKA substrate sequence results in increased efficacy of 480 nm-induced excitation and decreased efficacy of 400 nm-induced excitation. Thus, an increase in kinase activity is observed as an increase in the 488-/402-nm excitation ratio (Mehta et al., 2018). Excitation ratio imaging experiments were conducted 24 hours after transfection. Thirty minutes before imaging, cells were rinsed with serum-free medium and incubated in medium containing either vehicle (2.4 mg/ml HBC) or a cocktail of catecholamine transport inhibitors. The following transport inhibitors were used: corticosterone (500 μM) to inhibit OCT2 (IC_50_ = 500nM (Dirk Gründemann et al., 1998)), OCT3 (IC_50_ = 120 nM (Gründemann et al., 1998)) and PMAT (IC_50_ = 400 μM (Engel et al., 2004)); atomoxetine (10 μM) to inhibit NET; and GBR12909 (10 μM) to inhibit DAT. Dishes containing live astrocytes were then transferred to the temperature-controlled stage of a Nikon A1R confocal microscope to monitor excitation ratio responses to norepinephrine. Images at two excitation wavelengths (402 and 488 nm) were collected every 20 seconds for 3 minutes prior to, and for 30 min after, bath application of norepinephrine (final concentration 50 nM). This experiment was conducted on matched pairs (Groups: Vehicle and Transport Inhibitors) of dishes from 5 separate primary astrocyte culture preparations. Graphs were prepared using GraphPad Prism.

### Excitation Ratio Analysis

Nuclear fluorescence intensities for each excitation wavelength were normalized based on the average intensity during the baseline period for each nucleus. A ratio was then calculated by dividing the normalized intensity at 488 nm by the normalized intensity at 402 nm. Thus, increases in PKA are exhibited as increases in the ratio. Before examining group differences, any nuclei with excessive ratio drift during the baseline period were eliminated. The slope of the ratio during the baseline period was calculated for each nucleus, and the standard deviation of all slopes was calculated. Nuclei with baseline slopes exceeding two standard deviations of the mean were eliminated from subsequent analyses (one nucleus from the Transport Inhibitors group and two nuclei from the Vehicle group). For the remaining nuclei (n = 17 in the Transporter Inhibitors group and n=19 in the Vehicle group), average excitation ratios were calculated for the 3-minute baseline period and for each of three 10-min post-norepinephrine periods. The average excitation ratios for the two groups (Vehicle vs. Transport Inhibitors) were then analyzed with a mixed ANOVA across the 4 periods (Baseline vs. 0-10 vs. 10-20 vs. 20-30 minutes). Because we anticipated that the largest effect of the inhibitors would occur during the early post-norepinephrine periods, we conducted two planned interaction contrasts comparing the baseline periods with the first post-NE period (Baseline vs. 0-10 minutes); and comparing the first two post-NE periods (0-10 vs 10-20 minutes). Further comparisons were made using Tukey’s HSD.

## DATA AVAILABILITY

All data supporting the findings of this study are available from the corresponding author upon request.

## ACKNOWLEDGEMENTS

We would like to thank Colleen Lavin and Dr. Suresh Kumar for technical assistance with excitation ratio imaging.

## ADDITIONAL INFORMATION

### Funding

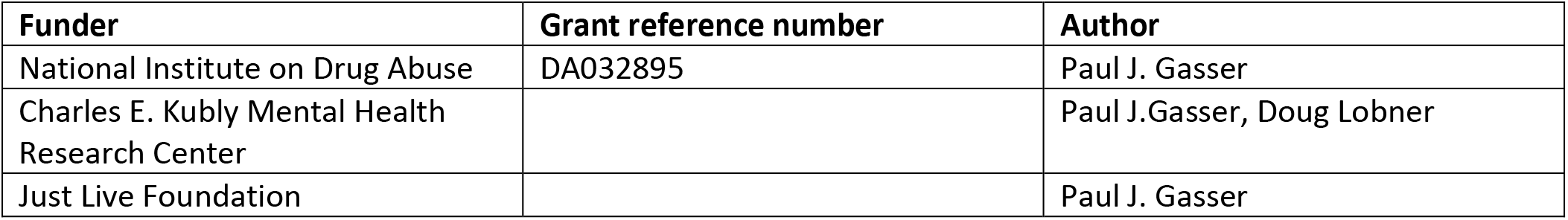

